# Longitudinal *in vivo* monitoring of axonal integrity after brain injury

**DOI:** 10.1101/2022.07.22.501178

**Authors:** Sergiy Chornyy, Julie A. Borovicka, Davina Patel, Min-Kyoo Shin, Edwin Vázquez-Rosa, Emiko Miller, Brigid Wilson, Andrew A. Pieper, Hod Dana

**Author notes:** Correspondence to: Hod Dana, Department of Neurosciences, Lerner Research Institute, Cleveland Clinic Foundation, 9500 Euclid Avenue, NC/30, Cleveland, OH 44195, USA, Andrew A. Pieper, Harrington Discovery Institute, University Hospitals Cleveland Medical Center, 11407 Euclid Ave, Cleveland, OH 44106, USA.

## Abstract

Traumatic brain injury-induced axonal degeneration leads to acute and chronic neuropsychiatric impairment, neuronal death, and accelerated neurodegenerative diseases of aging, including Alzheimer’s and Parkinson’s diseases. Thus, there is much interest in developing treatments that protect axons after injury. For this endeavor, extended comprehensive evaluation of axonal integrity in experimental systems is required to evaluate the efficacy of putative interventions in preclinical models. However, traditional histological tissue proccessing techniques are logistically prohibitive for assessments of long-term pathology. Here, we report a new method of longitudinally monitoring the functional activity of thalamocortical axons before and after injury *in vivo* in the same animal over an extended period of time. Specifically, we expressed an axonal-targeting genetically-encoded calcium indicator in the mouse dorsolateral geniculate nucleus and then recorded axonal activity patterns in the visual cortex in response to visual stimulation. We demonstrate the utility of this method for assessing *in vivo* aberrant axonal activity patterns after traumatic brain injury, as well as for evaluating the therapuetic efficacy of the neuroprotective P7C3-A20 pharmacologic agent *in vivo*. We found that P7C3-A20 treatment minimized most, but not all, of the pathological changes in axonal activity patterns after traumatic brain injury.

## Introduction

Worldwide, there are ~50 million annual cases of traumatic brain injury (TBI)^1^. These typically entail acute and chronic complications, including visual deficits, post-traumatic stress disorder, cognitive impairment, and major depression. TBI also incurs tremendous annual healthcare costs, reaching $13.1 billion in the United States alone, with an additional $64 billion in lost productivity^2^. An early and chronic consequence of TBI is axonal degeneration, which impairs brain circuitry and leads to neuronal death^3,4^. TBI also increases risk of aging-related neurodegenerative diseases, including Alzheimer’s disease (AD)^5^ and Parkinson’s disease^6^. Although the mechanism for this link is unknown, axonal degeneration is thought to play an important role^7^. To meet the urgent unmet need for neuroprotective strategies to stop axonal degeneration, it is critical to evaluate putative therapies in preclinical models. However, long-term assessments of axonal degeneration using traditional histological methods in animal models requires massive amounts of tissue for statistical power across a range of acute and chronic time points. Longitudinally monitoring axonal integrity *in vivo* in the same animals over extended periods of time, by contrast, could effectively evaluate putative neuroprotective agents more rapidly and with fewer animals.

Two-photon microscopy of the fluorescence signal from neurons expressing a genetically-encoded calcium indicator (GECI) allows *in vivo* monitoring of neuronal activity over multiple weeks^8^. Increases in intracellular calcium concentration following action potential firing can be detected in axons and used to measure activity patterns. For example, recording from axons of thalamocortical neurons projecting from the dorsolaterral geniculate neuclus (dLGN) to the primary visual cortex (V1) enables quantification of activity patterns tuned to different features of visual stimulation^9^. The development of an axon-targeting GECI (GCaMP6s-axon) further improves sensitivity through better signal-to-noise ratio^10^. Thus, we reasoned that nonlinear microscopy of axonal functionality after TBI could overcome some of the limitations of histology by generating data on pathology progression in the same animals over time. We applied this approach to identify that multimodal TBI affects the activity patterns of thalamocortical axons in the mouse visual system, and observed that this modulation may lead to changes in visual acuity. The NAD^+^-stabilizing neuroprotective compound P7C3-A20^11–17^ prevented most of these modulations. P7C3-A20 has previously shown potent neuroprotective efficacy in animal models of TBI^18–23^. Thus, efficacy in this model validates the sensitivity of the proposed method to identify neuroprotective effects, as well as revealing that thalamocortical projections may be protected after TBI.

## Materials and Methods

Surgical and experimental procedures were conducted in accordance with protocols approved by the Institutional Animal Care and Use Committee and Institutional Biosafety Committee of the Cleveland Clinic Lerner Research Institute, as well as the Louis Stokes Cleveland Veterans Affairs (VA) Medical Center Institutional Animal Care and Use Committee.

### Craniotomy surgeries and injection of viral particles

Nine C57BL/6J mice (5 males and 4 females, 3.5 months of age) were anesthetized using isoflurane (3% for induction, 1.5% during the surgery), placed on a heating pad, and then injected with local pain medication followed by exposure of the skull bone above the left hemisphere. A 3×5 mm^2^ craniotomy was drilled above the left cortex, and adeno-associated virus (AAV) was injected into the dLGN (9 injection spots per mouse at the following coordinates^9^: −1.8mm/−2.2mm (AP/LV), −2.2mm/−2.2mm, and −2.4mm/−2.2mm; injection depths were 2.7mm, 3mm, and 3.3mm). GcaMP6s-axon AAV1 solution (60 nl) was injected in each spot (Addgene catalog number 111262-AAV1). The craniotomy was covered with a custom-made glass window (Tower Optical Corporation) and cemented to the bone using dental cement (Contemporary ortho-Jet, Lang Dental). A custom-made metal head bar was attached to the skull. Mice were given post-operative care and 4 weeks for recovery to allow for sufficient GECI expression.

### Recording of axonal activity

Axonal activity recording was performed using a two-photon microscope with galvo-resonant scanners and GaAsP PMT detectors (Bergamo II, Thorlabs) at 950 nm excitation wavelength (Insight X3, Spectra-Physics) using a 16x, 0.8 NA objective (MRP07220, Nikon). Mice were lightly-anesthetized (0.5% isoflurane), injected with a sedative (Chlorprothixene Hydrochloride, Sigma; intra-muscular injection of 30 μl of 0.33 mg/ml solution), and placed on a heating pad in the dark^24–27^. Dense axonal labeling in Layer I of V1 was apparent four weeks after AAV injection. Visual stimulation was presented to the right eye, generated using the psychophysical toolbox^28,29^ in Matlab (Mathworks) on an LCD monitor (30×36 cm^2^ display, located 15 cm in from of the mouse right eye, tilted 45 degrees with respect to the nose line, and covered with a blue plexiglass to minimize fluorescence signal contamination) that subtended an angle of ±50° horizontally and ±45° vertically. The visual stimulus consisted of a drifting grating moving in 1 of 8 directions for 4 sec, followed by 4 sec of gray display. This stimulation cycle was repeated 5 times. Fields of view (FOVs) of ~200×200 μm^2^ with 512×512 pixels at 30Hz were recorded. Typically, 5 different FOVs were recorded from each mouse in each recording session. The different FOVs were identified by their location with respect to Bregma (using a motorized stage with 1-μm resolution), as well as the shape of the local vasculature. During subsequent recordings, we returned to the same cooridnates and used the local vasculature as a map (See Supplementary Video 1) to locate the brightest axons within the FOV so that we could then record from the same location. If the exact location of a secific FOV could not be determined due to changes in the brain condition (i.e., degeneration of axons or changes in vasculature shape after TBI), then we recorded from a nearby location within several hundred microns from the original FOV. Over the following weeks, we continued recording from the same modified location. FOVs were excluded from analysis if there were severe movements, or if visual stimulation was not synchronized with recording.

### Multimodal TBI

Mice were anesthetized with ketamine/xylazine (100/10 mg/kg) via intraperitoneal injection and placed in an enclosed chamber constructed from an air tank partitioned into two sides and separated by a port covered by a mylar membrane. Pressure in the side not containing the mouse was increased to cause membrane rupture at 20 pounds per square inch (PSI), generating ~1.0- to 1.5-m/s air jet flow of 137.9 ± 2.09 kPa that passes through the animal’s head. The head was untethered in a padded holder while the body was shielded by a metal tube. The jet of air produced upon membrane rupture produces a collimated high-speed jet flow with dynamic pressure that delivers a compressive impulse. Variable rupture dynamics of the diaphragm through which the jet flow originates also generate a weak shock front. There is also a minor component of acceleration– deceleration injury to the unrestrained head. The sham injury (control) group was anesthetized and passed through the same process except for the injury. After several hours of monitoring, mice were returned to regular housing. All mice were included in the study and were randomly assigned to experimental groups; no experimental blinding was used.

### P7C3-A20 treatment

P7C3-A20 was dissolved in 2.5% vol of dimethyl sulfoxide, followed by addition of 10% vol Kolliphor (Sigma-Aldrich) and vigorous vortexing. This solution was diluted in 87.5% vol of D5W (filtered solution of 5% dextrose in water, pH=7.0). Mice in the P7C3-A20 treatment group were intraperitoneally injected with 10mg/kg P7C3-A20 every morning throughout the recording period, starting one day after TBI. The other two groups received no treatment.

### Data analyses

Small drifts and movements of the imaged area throughout recording were corrected using TurboReg plug-in of ImageJ^30^. Axonal segmentation was done using Suite2P^31^ with experimenter supervision. Remaining analyses were conducted with custom Matlab scripts. Fluorescence signal was averaged from all pixels within each segment. Neuropil signal was subtracted, as previously done^26,32^, with a coefficient of 0.7. Baseline fluorescence (F_base_) for each segment was calculated by averaging fluorescence for 0.66 sec before visual stimulation. The response amplitude (F_resp_) was calculated as peak amplitude of fluorescence signal during the drifting grating appearance and calculating ΔF/F_0_=(F_resp_-F_base_)/ F_base_.

Detection of tuning of segments to visual stimulus (tuned segments) was calculated as in previous studies^25,26,32^ using a one-way ANOVA test (p=0.01) between mean fluorescence signal during stimulus presentation vs. during presentation of gray display. Among the tuned segments, a second one-way ANOVA test (p=0.01) between mean fluorescence signal during presentation of each of the 8 directions determined whether these segments had an orientation preference (oriented segment). Orientation sensitivity index (OSI) and direction sensitivity index (DSI) were calculated as previously defined^32,33^. The preferred angle for each tuned segment was determined as the grating angle that elicited the highest fluorescence amplitude change (θ_pref_). We calculated OSI=(θ_pref_-θ_ortho_) /(θ_pref_+ θ_ortho_), where θ_ortho_= θ_pref_+π/2. Similarly, DSI=(θ_pref_-θ_opposite_) /(θ_pref_+θ_opposite_), where θ_opposite_= θ_pref_+θ.

### Statistical analyses

For comparing long-term changes, we grouped together data from 2-3 recording sessions (pre-TBI, weeks 1-2, 3-4, and 5-7 post-TBI). To assess the effects of time, treatment group, and their interaction given the repeated measures of FOV and multiple FOV measured within each mouse, we estimated linear mixed-effects models considering FOV within mouse as nested random effects. The significance of the modeled fixed effects of time, treatment group, and their interaction was assessed with F-tests using the marginal sum of squares and considered significant at p < 0.05. Model assumptions were assessed for each endpoint and square-root transformations were imposed for percentage of tuned and oriented segments, response amplitude, OSI, and DSI. Model-estimated marginal means and 95% confidence intervals were calculated and overlayed on observed data in figures with distinct line types or symbols for each mouse within a treatment group.

When a significant interaction was detected, post-hoc tests were implemented to identify which paired treatment groups differed in any of the four time estimates (joint hypothesis tests of time parameters assessed as F-tests), or in which treatment groups we observed significant changes over time (pairwise tests of time parameters within treatment group as t-tests with containment degrees-of-freedom). Within each endpoint, the p-values of multiple post-hoc tests were adjusted using the Holm method and considered significant at p < 0.05. All analyses were performed in R (Version 4.1.3)^34^ and included functions from the nlme and emmeans packages^35,36^.

### Data availability

The data that support the findings of this study are available from the corresponding authors upon reasonable request.

## Results

### Monitoring axonal activity before and after multimodal TBI

All mice were injected with AAV solution carrying the GCaMP6s-axon GECI sequence^10,26^ into their dLGN. Weekly recording sessions started four weeks later in mice that were lightly anesthetized using a combination of chorprothixene hydrochloride and isoflurane, as previously used to characterize functional responses of primary visual cortex neurons^24–27,32,37^. Functional activity of dLGN axonal projections in Layer I of V1 was monitored before and after TBI (Fig. 1A-B), and as a function of treatment with P7C3-A20, a neuroprotective agent that blocks axonal degeneration by restoring cellular levels of NAD^+^ after TBI^19,22^. Presenting a multi-directional drifting grating movie enabled the assessment of dLGN axons’ visual properties^9,33^ as a function of exposure to TBI and treatment. In this TBI model, an overpressure chamber delivers global concussion, acceleration/deceleration injury, and early blast wave exposure in an isolated manner to the head of an anesthetized mouse. This generates a reproducible pattern of axonal degeneration, cognitive impairment, blood-brain barrier disruption, systemic metabolic alterations, and blood biomarkers that reliably mimic human TBI^38^. Following 2-3 baseline (pre-TBI) axonal activity recordings, 9 mice were randomly assigned to three groups: 1) TBI, 2) TBI treatment, in which mice received TBI and daily P7C3-A20 starting 24 hours after injury, and 3) sham injury (control), in which mice were subjected to sham-TBI. Post-TBI recordings were conducted once a week over seven weeks for all groups, starting three days after TBI (Supp. Fig 1).

**Figure 1.**
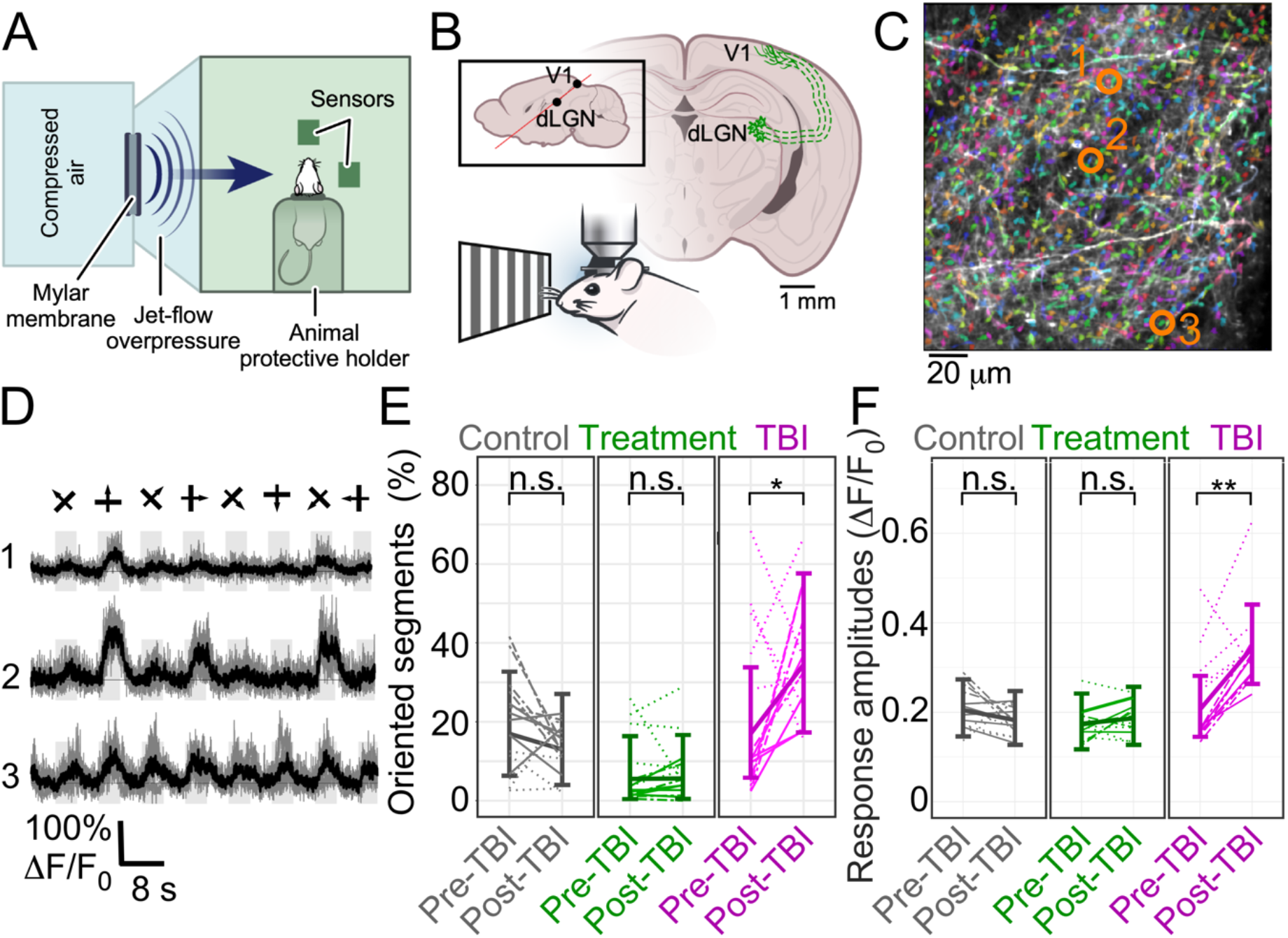
Multimodal TBI causes short-term changes in axonal properties, which are minimized by P7C3-A20 treatment. **(A)**. Schematic of the overpressure chamber used to produce multimodal TBI in mice. (**B)**. Illustration of the experimental setup. Mice were injected with GCaMP6s-axon into their dLGN, and axonal activity from Layer I of V1 was recorded during the presentation of a drifting grating movie, both before and after TBI. (**C)**. Example image of axons labeled with GCaMP6s-axon (white over black background) overlaid with the segmentation from Suite2P. Segment colors were randomly selected to emphasize the different segments in the image. (**D)**. Fluorescence traces of 3 example segments (indicated by orange circles in **C**). Single trials (gray) and averages of five trials (black) are overlaid. Eight grating motion directions are indicated by arrows as shown above the traces. (**E)**. The model-estimated mean fraction of axonal segments with significant orientation preference (ANOVA, p<0.01) was increased from 16.9% to 34.5% for the TBI group between the last weekly recording before injury and the first recording 3-5 days following the injury. This fraction remained similar in the sham control group (gray) and P7C3-A20 treatment group (green) (P=0.004, F-test, for group-time interactions; post-hoc time comparisons within treatment group found a significant Holm-adjusted increase of P=0.047 for the TBI group, but not for the other groups). *, p<0.05; n.s., not significant). (**F)**. Fluorescence response amplitudes to visual stimulation showed a significant group-time interaction (P<0.001) with a significant increase in the model-estimated mean for the TBI group of 67% from their pre-TBI level (P=0.002, Holm-adjusted), while the treatment and control groups showed no significant changes (same FOVs as in **E;** **, p<0.01). Within each experimental group, different lines (solid, dotted, and dash-dotted) show data from different mice. The thick, solid lines connect the model-estimated mean values and the error bars show the model-estimated 95% confidence intervals. Figures and models each include 88 measures of 46 FOVs in 9 mice (3 per experimental group).

### Changes in axonal activity patterns following TBI and the protective effect of P7C3

We detected hundreds of axonal segments within each 200×200 μm^2^ FOV (median of 624 segments/FOV, range 240-1196, total number of 402 recordings from 58 FOVs in 9 mice that were segmented using Suite2P^31^; Fig. 1C). Individual segments exhibited orientation- and direction-tuned activity when a drifting grating stimulation of eight movement directions was presented to the contralateral eye^9^ (Fig. 1D, see Materials and Methods). Response properties were substantially increased in the TBI group between the last weekly recording before TBI and the first recording after TBI, compared to the sham control group. The model-estimated mean fractions of axonal segments significantly tuned to the stimulus and showing orientation selectivity increased by 74% and 104%, respectively (Fig. 1E, Supp. Figs. 2-3, and Supp. Video 1). Blocking axonal degeneration with P7C3-A20 prevented these TBI-induced changes, and no significant changes from pre- to post-TBI were detected in the treatment group. Axonal activity level during stimulation, measured as fractional increase in GECI signal, was also increased from model-estimated mean [95% CI] of 20.8% [14.5%, 28.1%] to 34.7% [26.3%, 44.1%] after TBI, and was not significantly changed in the control or treatment groups (20.5% [14.6%, 27.3%] vs. 18.2% [12.7%, 24.7%] and 17.4% [11.7%, 24.2%] vs. 18.6% [12.7%, 25.7%]; mean [95% CI]; Fig. 1F). Thus, TBI caused significant short-term increases in axonal activity that were prevented by treatment with P7C3-A20.

Over the seven weeks following TBI, the fraction of axonal segments with orientation preference showed significant interactions between the experimental groups over time (P=0.017, F-test). The TBI group remained above its pre-TBI levels during the entire recording period, with percentage point increases in the model-estimated mean of 15%, 11%, and 15% on weeks 1-2, 3-4, and 5-7, respectively (Fig. 2A). The treatment group showed more modest increases of 4%, 6%, and 6% on weeks 1-2, 3-4, and 5-7, respectively, and the sham control showed a slight change compared to the baseline values. Comparing the effects of time between pairs of experimental groups, the TBI group differed significantly from the control group (Holm-adjusted P=0.027), but not the other pairs (Fig. 2A). The fraction of tuned segments showed a similar trend to the oriented segments (Supp. Fig. 4A) and the ratio of oriented-to-tuned segments gradually increased for all groups, but without a significant interaction of time or treatment (P=0.052 and P=0.059, respectively; F-test). This ratio for the TBI group increased from a model-estimated mean value of 46.3% before TBI to 56.4%, 63.8%, and 65.6% in weeks 1-2, 3-4, and 5-7, respectively. The treatment group showed delayed and smaller increases from 31.9% before TBI to 47.9% in weeks 5-7, and the sham control group showed no specific trend (Supp. Fig. 4B). Interestingly, we also found a non-significant reduction of 15.3% in the model-estimated mean number of detectable axonal segments across all weekly-recorded FOVs in the TBI group, which is in line with reported axonal degeneration in this injury model^3^ (P=0.375, F-test; Supp. Fig. 5).

**Figure 2.**
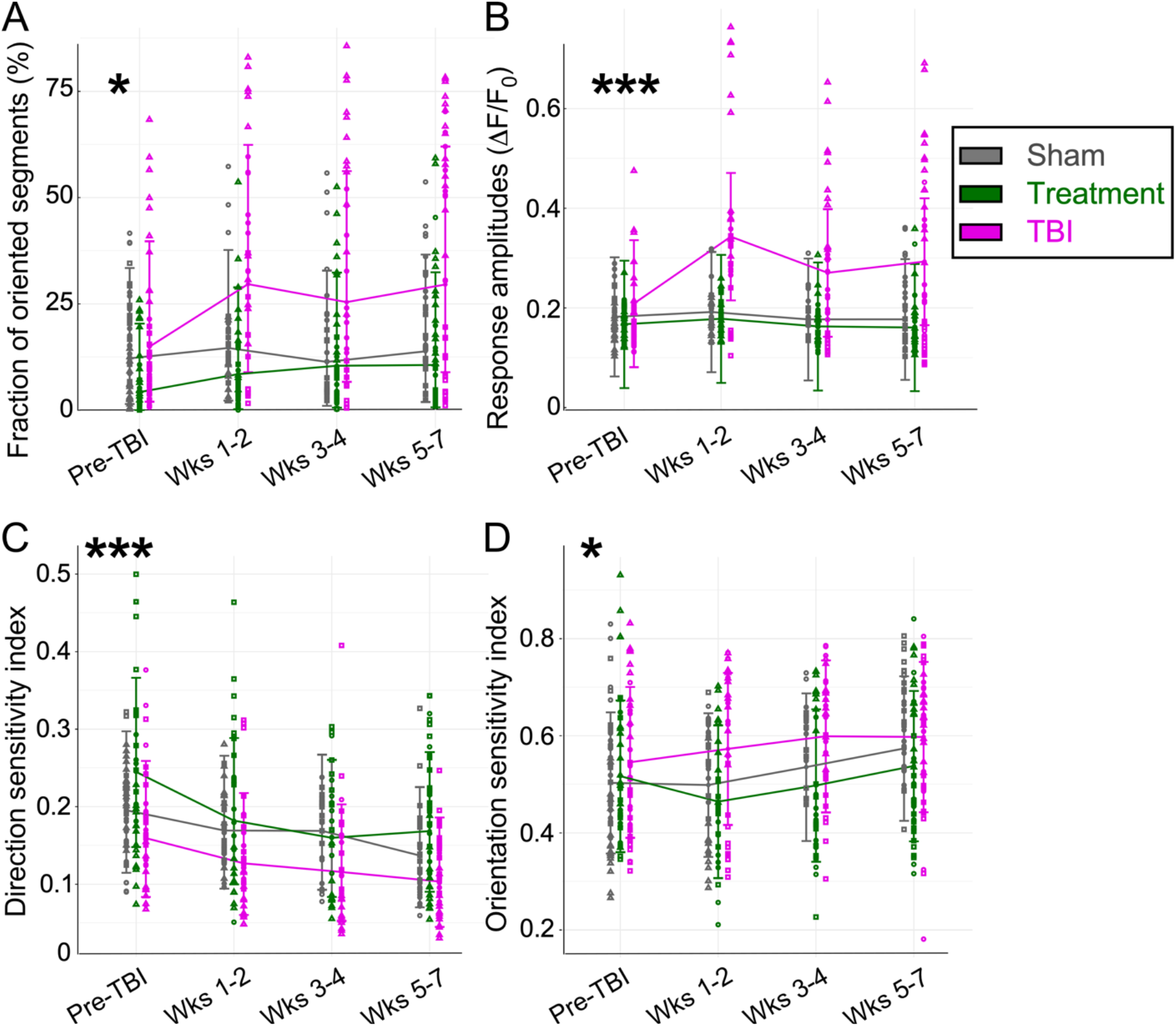
Multimodal TBI causes long-term deficits to thalamocortical axonal activity patterns, tuning, and acuity, which are mostly reversed by P7C3-A20 treatment. (**A)**. The fraction of oriented segments showed a significant group-time interaction (P=0.017, F-test) with significant differences in the effects of time for the TBI group compared to the sham control group (P=0.027, Holm-adjusted), but not for the P7C3-A20 treatment group (P=0.173, Holm-adjusted). (**B)**. The measured fluorescence response amplitudes for a drifting grating simulation showed a group-time interaction (P<0.0001, F-test) and model estimated mean amplitudes were 63% higher in TBI mice in the first 1-2 weeks after TBI compared to the pre-TBI period. Response amplitudes remained higher by 29% and 35% on weeks 3-4 and 5-7, respectively. The effects of time differed significantly between the TBI group and the sham control group as well as between the TBI and treatment groups (P<0.0001 for both comparisons, Holm-adjusted). The sham control and treatment groups did not show such an increase, and there was no significant difference in time effects between them (P=0.999, Holm-adjusted). (**C)**. The median DSI value of the tuned segments showed a significant decrease during 7 weeks of post-TBI recording (P<0.0001, F-test), but without a group-time interaction (P=0.084, F-test). The decreases for the TBI group were greatest between pre-TBI and weeks 1-2 and continued throughout the following recordings. The sham control group showed a more gradual decrease, and the decrease in the treatment group was greatest between pre-TBI and weeks 1-2, but ceased after weeks 3-4. **(D)**. The median OSI value of the tuned segments showed a significant increase during 7 weeks of post-TBI recording for all groups (P=0.025, F-test), but without a group-time interaction (P=0.257, F-test). The TBI group showed a larger increase within the first 1-2 weeks after injury compared to the other groups (same FOVs as in **C**). Within each group, data from individual mice are represented by different markers (square, triangle, circle). The lines connect the model-estimated mean values and the error bars show the model-estimated 95% confidence intervals. Figures and models each include 402 measures of 58 FOVs in 9 mice (3 per treatment group).

The ability of this axonal-monitoring-based method to detected different effects of TBI on the neuronal circuitry was demonstrated by monitoring individual mice within the experimental groups. For example, while all TBI mice exhibited enhanced axonal response amplitudes in the first post-TBI week, one mouse in this group also showed subsequently reduced response amplitudes and tuning metrics over the following weeks, whereas the other two showed increased response amplitudes and tuning (Supp. Figs. 6-7). We quantified the change in median response amplitudes over time and found a significant group-time interaction (P<0.0001). The increase over time in the TBI group was significantly different from both the treatment and sham control groups (P<0.0001 for both, Holm-adjusted for multiple comparisons), and the treatment and sham control groups showed no significant difference across time (adjusted P=0.999). Interestingly, most of the increased response amplitudes for the TBI group occurred within the first two weeks after injury and remained at the same high level thereafter (Fig. 2B). Taken together, our results show that *in vivo* monitoring of axonal tuning after TBI provides a real-time indicator of axonal degeneration and post-TBI pathologic or neuroprotective changes to neuronal circuitry.

### Changes in visual acuity over time

Finally, we assessed visual acuity by calculating the DSI and OSI for individual tuned segments^9,33^ (Fig. 2C-D; see Materials and Methods). We identified a significant decrease in DSI for all groups over time (P<0.0001, F-test) and a significant increase in the OSI over time (P=0.0247, F-test). For the sham control group, we observed a ~30% longitudinal decrease in model-estimated mean DSI and a ~14% increase in OSI during the recording period, which may be related to physiological changes in the visual system during its maturation^39^. For both DSI and OSI, there were no significant interactions between time and experimental groups (P=0.084 and P=0.257, respectively). The TBI group exhibited similar trends to the sham control group, but more quickly. Model-estimated mean DSI medians significantly decreased by 20% within the first 2 weeks and maintained similar levels thereafter. Similarly, model-estimated mean OSI increased by 5% on weeks 1-2 post-TBI and maintained similar levels over the following weeks. The treatment group showed similar DSI changes compared to the other groups, decreasing ~31% during the recording period, while its model-estimated mean OSI showed no significant trends. Therefore, all groups showed modulated visual acuity over time, and both the TBI and treatment groups exhibited different patterns compared to the sham control group, indicating possible persistence of some TBI-related deficits.

## Discussion

Here we report a new method for *in vivo* monitoring of axonal integrity that enables longitudinal quantification of TBI-induced acute and chronic axonal damage. Two-photon microscopy of GECI signal offers several advantages over histiology-based methods, beacuse it allows repeated measurements from the same brain region within the same mouse. This facilitates quantitative assessments of pathology progression within the same subject. Moreover, functional imaging of axonal fluorescence changes to visual stimuli enables the assessment of mouse visual system properties^33^. TBI-induced axonal pathology was detected within days after injury and did not return to baseline without treatment. Treatment with P7C3-A20^19,22^ eliminated most, but not all, of the identified pathologies (Fig. 1E-F, Fig. 2, Supp. Figs. 2, 4, and 5). Thus, this new method allows *in vivo* identification of variability across animals (Supp. Fig. 6-7) and quantification of the neuroprotective efficacy of pharmacologic treatment after brain injury. However, we note that comprehensive description of variability across mice in this *in vivo* monitoring technique after TBI will require testing a substantially larger cohort size. Here, we limited our scope to the proof-of-principle of this new technique. Depending upon the injury or disease paradigm, application of this technique will require different numbers of mice to achieve appropriate statistical power.

Axonal degeneration plays a central role in multiple neurodegenerative conditions, and the small size and large distribution of axons across multiple brain regions makes it challenging to estimate their condition using conventional tissue histological processing. Moreover, postmortem histology precludes monitoring of trends of recovery or degeneration of axons within the same animal. Repeated measurements of axonal function within the same animal allow reducing the between-animal variability that is inevitable when acute axonal assessment methods are used. In addition, within-subject monitoring increases the confidence in assessing the time course of recovery and pathology reversal due to treatment compared to studies in which different animals are euthanized at various time points. Therefore, our new method may assist in reducing the required number of animals for detecting several key pathological changes after TBI and will facilitate more efficient testing of new treatments. This new approach can be similarly applied to monitor axonal degeneration in other preclinical models of injury or neurodegeneration, in order to investigate pathology and to facililtate evaluation of putative neuroprotective treatments for these conditions.

## Abbreviations

TBI: traumatic brain injury
AD: Alzheimer’s disease
dLGN: dorsolatreral geniculate neuclus
GECI: genetically-encoded calcium indicator
V1: primary visual cortex
FOV: field of view
DSI: direction selectivity index
OSI: orientation selectivity index
AAV: adeno-associated virus
AP: anterior-posterior
LV: lateral-ventral
PSI: pounds per square inch

## Acknowledgments

We thank Drs. Christopher Nelson and Dimitrios Davalos for reading and commenting on the content of this manuscript. We thank the HHMI Janelia GENIE project for sharing the GCaMP6s GECI, and Dr. Lin Tian’s lab for sharing the GCaMP6s-axon construct. We thank Dr. Marius Pachitariu and his colleagues for sharing the Suite2P analysis platform.

