## Supplementary material for "Longitudinal *in vivo* monitoring of axonal integrity after brain injury": Supp. Fig.

### Longitudinal *in vivo* monitoring of neuroprotection after traumatic brain injury

#### Supplementary information

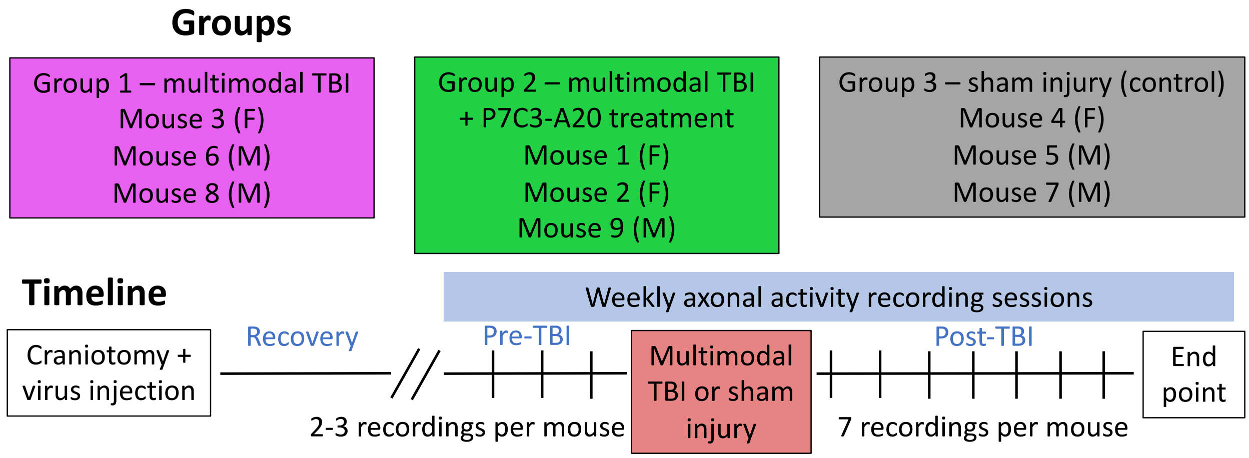


**Supplementary Figure 1. Experimental design and timeline.** Nine male and female mice (M and F, respectively) were randomly divided into 3 groups: TBI, TBI and P7C3-A20 treatment, and sham injury (control). All mice were implanted with cranial windows and injected with GCaMP6s-axon into their dLGN. We recorded 2-3 sessions of baseline visual stimulated activity before exposing mice in the TBI and treatment groups to multimodal TBI. Mice in the sham injury group were anesthetized and placed inside the multimodal TBI device without activating the overpressure chamber. We continued recording axonal activity for seven weeks after injury, during which time the treatment group was administered 10 mg/kg of P7C3-A20 daily.

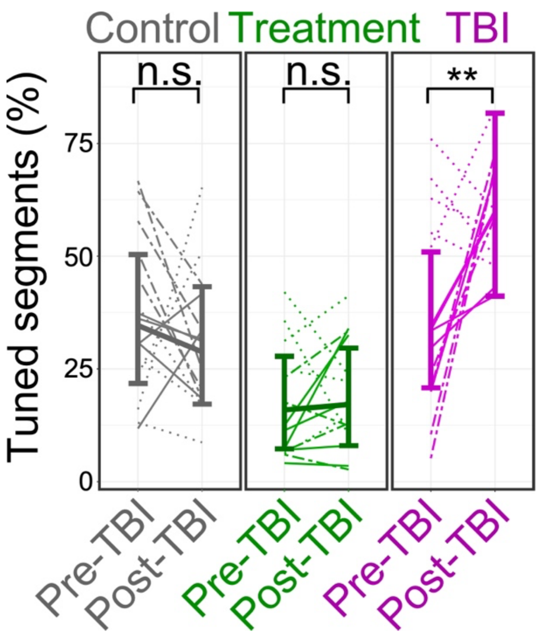


**Supplementary Figure 2. The short-term increase in axonal tuning after multimodal TBI is blocked by P7C3-A20 treatment.** The mean fraction of axonal segments that were significantly tuned ($P<0.01$, ANOVA test) to the presented visual stimulus showed a significant difference between the experimental groups ($P=0.005$, F-test). For the TBI group, the fraction was significantly increased by 74% (magenta; $P=0.002$, with Holm adjustment for multiple comparisons) in the days before and after TBI, but not for the sham control group (gray; $P=0.595$). P7C3-A20 treatment protected against this increase (green; $P=0.769$); **, $p<0.01$; n.s., not significant). For each experimental group, different lines (solid, dotted, dash-dotted) show data from different mice, error bars show the 95% confidence interval, and the model mean values are connected with lines.

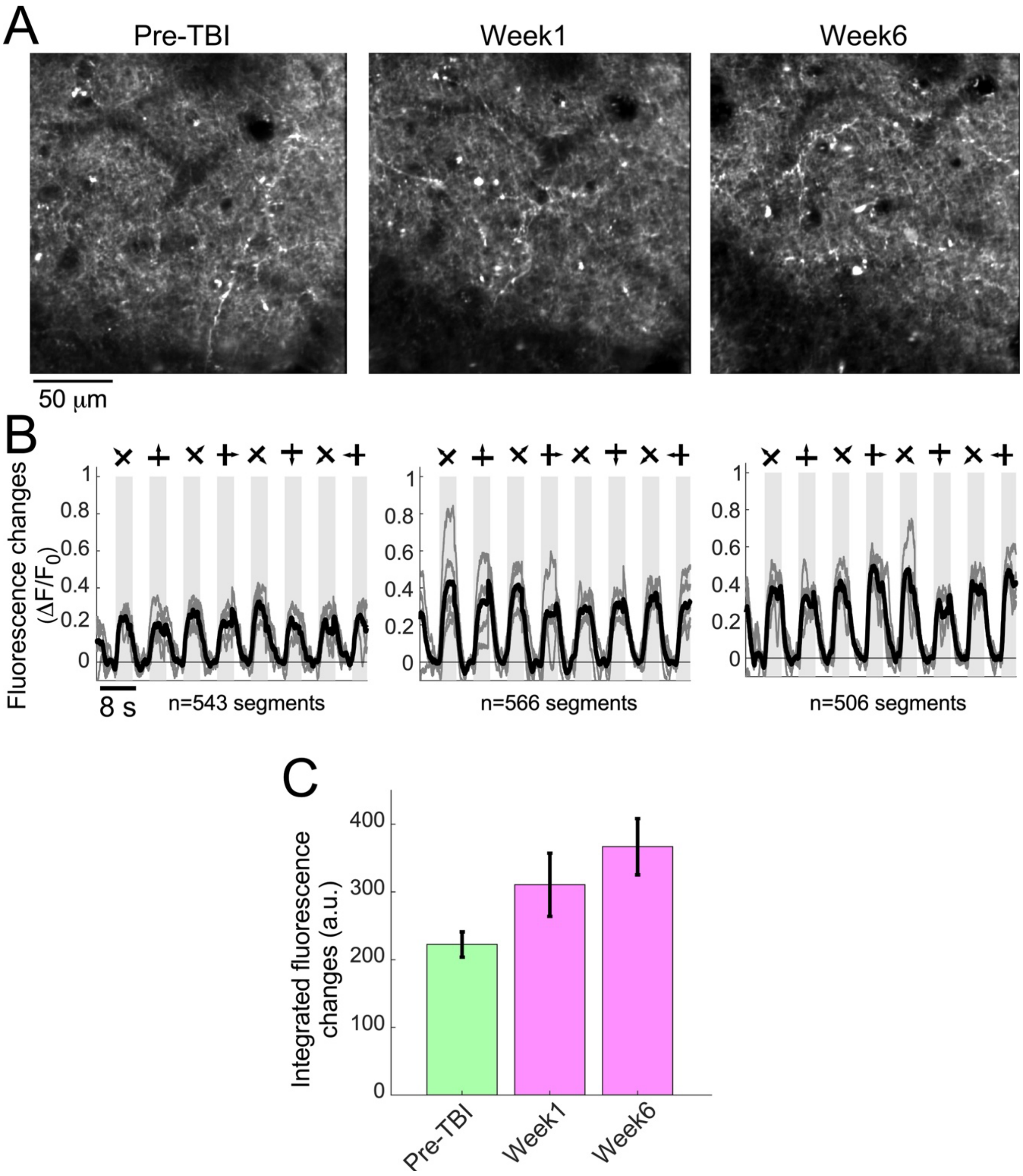


**Supplementary Figure 3. Repeated recordings from the same FOV. A.** Example mean images of the same FOV (mouse 8, TBI group) recorded 1 day before TBI (left), 4 days after TBI (middle), and 6 weeks after TBI (right). Activity movies from these recordings are shown in Supplementary Video 1. **B.** Mean $\Delta F/F_0$ traces across all detected segments for each FOV shown in **A**. Single trials (gray) and averages of five trials (black) are overlaid. Eight grating motion directions are indicated by arrows as shown above the traces. **C**. Sum of the $\Delta F/F_0$ traces in **B** shows an increase in response amplitudes after TBI (mean±s.d. for the 5 trials).

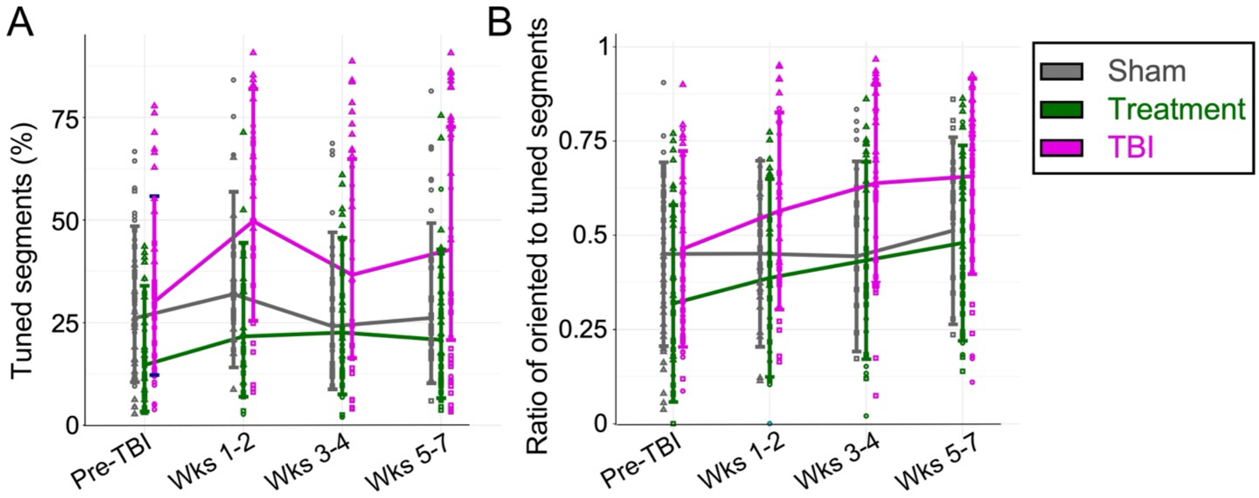


**Supplementary Figure 4. Increase in the fraction of tuned segments and the oriented-to-tuned segment ratio for the TBI group. A.** The percentage of tuned segments (P<0.01, one-way ANOVA) was increased for the TBI group over the weeks after injury, this increase was smaller the treatment group, and no increase was identified for the sham control, although no significant interaction of time and groups was detected (P=0.052, F-test). **B.** The ratio of oriented segments to tuned segments was increased for the TBI and treatment groups, but not the sham control group, although no significant interaction of group and time was detected (P=0.059, F-test). For each group, data from single mice are represented by different markers (square, triangle, circle), the error bars show the 95% confidence intervals, and the model-estimated means are connected with lines.

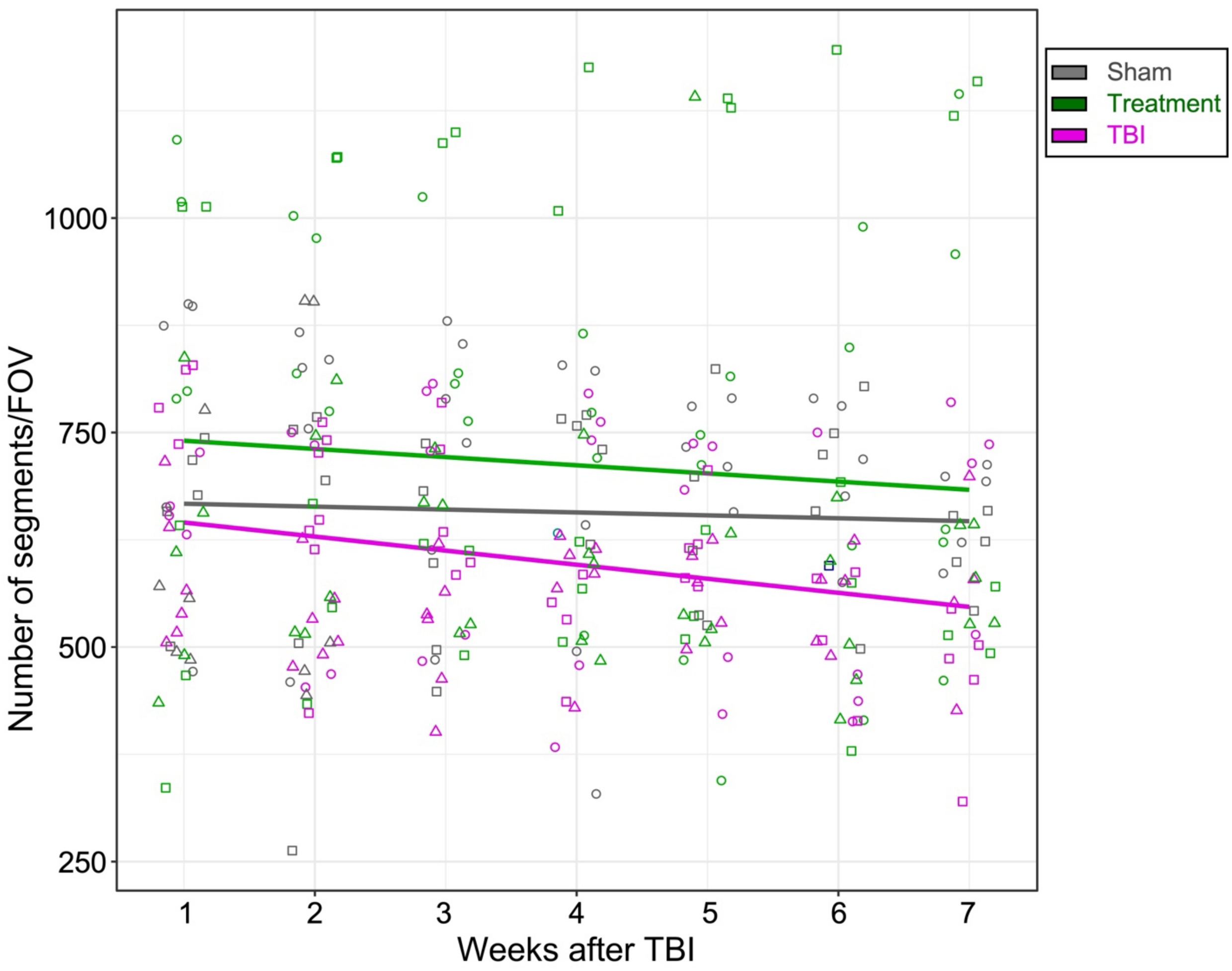


**Supplementary Figure 5. Gradual decrease in the number of detected segments after TBI.** The number of segments per FOV showed a decrease in the TBI group (magenta), with a model-estimated decrease of 15.3% from Week 1 to Week 7. Smaller decreases were identified in the sham control (gray) and treatment (green) groups. These changes are consistent with reported findings on axonal degeneration in this multimodal TBI model. We note that the changes in our study were not significant for time, treatment group, or their interaction (P=0.375, F-test), suggesting that either more mice per group and/or longer recording periods are required to identify significant differences. For each group, data from single mice are represented by different markers (square, triangle, circle), and the model means are connected with lines.

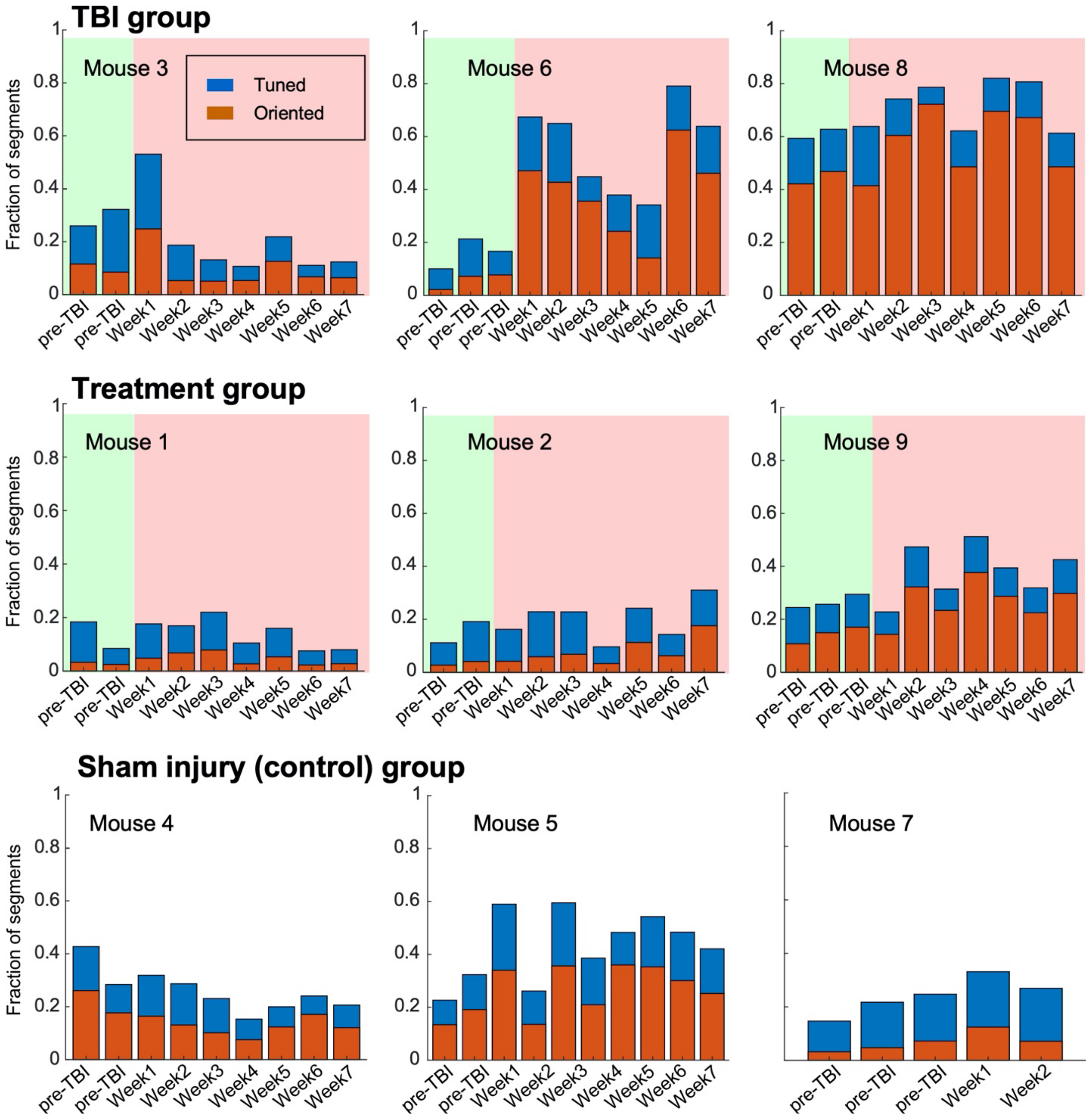


**Supplementary Figure 6. Tuning and orientation preferences of axonal segments from individual mice before and after multimodal TBI.** Longitudinal monitoring of the fractions of tuned and oriented segments for individual mice. The fractions of tuned and oriented segments (blue and orange bars, respectively) from all recorded segments are shown for the multimodal TBI group (upper row), TBI and treatment group (middle row), and sham injury group (bottom row). Green and red backgrounds indicate the pre-TBI and post-TBI periods, respectively. Data from all recorded FOVs on the same date were compiled to calculate the respective fractions. Note that data from mouse 7 was acquired for only 5 recording dates.

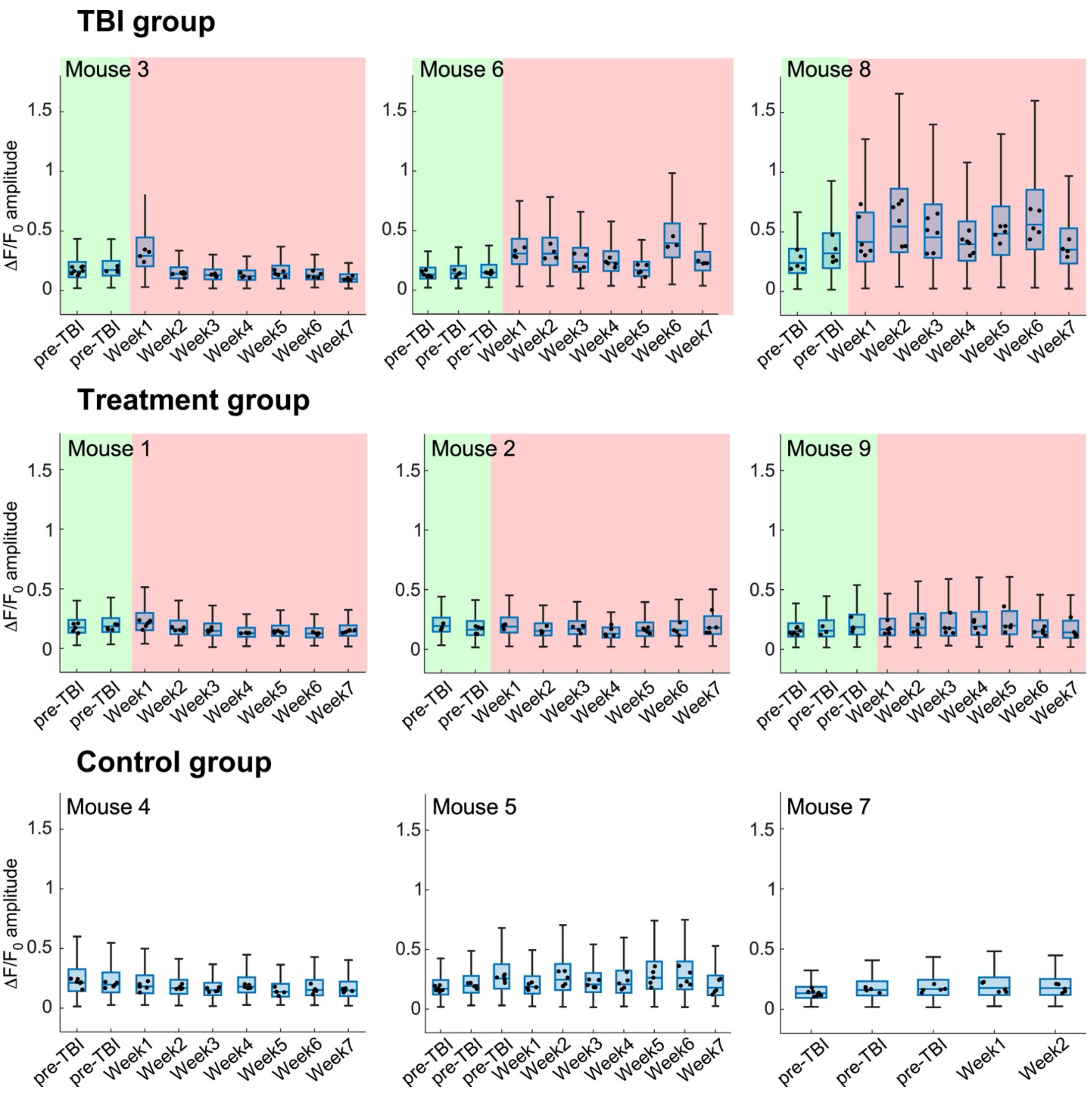


**Supplementary Figure 7. Longitudinal monitoring of the response amplitudes for individual mice before and after multimodal TBI.** Data are shown for the multimodal TBI (upper row), treatment (middle row), and sham control (bottom row) groups. Each box represents the 25-75 percentile range of $\Delta F/F_0$ amplitude from all recorded segments on a specific recording session. The whisker spans the smallest among the range of the entire data set, or 1.5 times the interquartile range. For each recording date, the median values from all recorded FOVs are overlaid on the boxplot data (black dots).

**Supplementary Video 1. Repeated recordings from the same FOV.** Activity movies from the same FOV, as described in Supp. Fig. 3, are shown. All movies are synchronized, and the timestamp shows the recording times for all FOVs. The appearance of an arrowhead in the upper-left corner indicates the appearance and movement direction of the drifting grating stimulus for all recordings. Raw recording data were smoothed using a 9-frame running average window and are displayed 3x faster than real time (90 frames per second).
